# Autism_genepheno: Text mining of gene-phenotype associations reveals new phenotypic profiles of autism-associated genes

**DOI:** 10.1101/2021.03.24.436848

**Authors:** Sijie Li, Ziqi Guo, Jacob B. Ioffe, Yunfei Hu, Yi Zhen, Xin Zhou

## Abstract

Autism is a spectrum disorder with wide variation in type and severity of symptoms. Understanding gene–phenotype associations is vital to unravel the disease mechanisms and advance its diagnosis and treatment. To date, several databases have stored a large portion of gene–phenotype associations which are mainly obtained from genetic experiments. However, a large proportion of gene–phenotype associations are still buried in the autism-related literature and there are limited resources to investigate autism-associated gene-phenotype associations. Given the abundance of the autism-related literature, we were thus motivated to develop Autism_genepheno, a text mining pipeline to identify sentence-level mentions of autism-associated genes and phenotypes in literature through natural language processing methods. We have generated a comprehensive database of gene-phenotype associations in the last five years’ autism-related literature that can be easily updated as new literature becomes available. We have evaluated our pipeline through several different approaches, and we are able to rank and select top autism-associated genes through their unique and wide spectrum of phenotypic profiles, which could provide a unique resource for the diagnosis and treatment of autism. The data resources and the Autism_genpheno pipeline are available at: https://github.com/maiziezhoulab/Autism_genepheno.

## Introduction

Many human diseases are the result of a complex interplay between genotype, the set of genes that each organism carries, and the environment. Gene–phenotype association analysis plays an important role in understanding the mechanisms of different genetic diseases, however substantial gaps in our knowledge remain^1–4^. Autism Spectrum Disorder (ASD) is such a genetic condition with heritability estimated at close to 90%^5^. Despite many Genome-Wide Association Studies (GWAS), known genetic effects can only account for only 24–33% of cases. For the majority of patients, the etiology is unknown. Part of the problem is that autism is a “spectrum” disorder, with wide variation in the type and severity of symptoms patients have experienced^6, 7^. To date, the SFARI autism database has curated close to 1,000 genes associated with autism, and this number is increasing every year^8^. A vast amount of gene–phenotype associations has been reported in the biomedical literature. Understanding how ASD phenotypes rise from a patient’s genetic composition is highly valuable to advance diagnosis and treatment^4, 9–14^.

Given the complexity of the condition, the number of genes implicated, and the variability of symptoms in individual patients, there is an enormous amount of literature for Autism studies. Phenotypes have been made available in several databases such as ClinVar^15^, Online Mendelian Inheritance in Men (OMIM)^16^, and the Human Phenotype Ontology (HPO)^17^ by many genetics experiments. Phenotype ontologies have also been developed to easily integrate and compare phenotypes among different species. There is also a recent integration of phenotype ontologies for autism: Autism Spectrum Disorder Phenotype Ontology (ASDPTO)^18^. Systematic extraction of gene-phenotype association from this literature is challenging. Development of automated text mining tools to extract a comprehensive database for gene–phenotype associations is therefore highly valuable and can alleviate the burden of manual curation^2, 19^.

We were thus motivated to develop Autism_genepheno, a text mining pipeline to generate a comprehensive database of gene-phenotype associations in ASD that can be easily updated as new literature becomes available. We designed Autism_genepheno to identify sentence-level mentions of autism-associated genes and phenotypes from HPO and ASDPTO in literature through natural language processing (NLP) methods. To further determine the strength of gene-phenotype associations, we applied a measure of Normalized Pointwise Mutual Information (NPMI) on the comprehensive extracted gene-phenotype associations. We used Autism_genepheno to process the last five years of autism publications and generated a large gene-phenotype association matrix for autism-associated genes classification. We evaluated gene-phenotype associations from Autism_genepheno through comparison with existing gene-phenotype associations in reference databases. Gene Ontology (GO) analysis we performed for some specific clusters of autism-associated genes further confirmed the strength of our gene-phenotype associations. The gene classification analysis for SFARI genes also revealed specific phenotypic profiles for different classes of SFARI genes which conferred autism risk at different levels, and we were able to select the top 10% SFARI genes with the most influential roles in a genetic interaction network based on their phenotypic profiles.

## Methods

### Pipeline Overview

We assumed a gene-phenotype association exists if gene-phenotype co-occurrence pairs were detected in the same sentence of the published articles. We introduced a pipeline to detect gene-phenotype associations of Autism Spectrum Disorder from PubMed Central (PMC) articles. The overview pipeline is shown in Figure 1. The input of the pipeline is a corpus consisting of autism research articles in XML. The output is a list of quantitative gene-phenotype associations detected from the corpus. The pipeline includes three steps: 1) detecting sentence-level gene-phenotype associations with curated gene list and phenotype list; 2) standardizing the detected phenotypes and their top-level phenotypic category with HPO; and 3) ranking gene-phenotype associations by Normalized Pointwise Mutual Information (NPMI).

**Figure 1.**
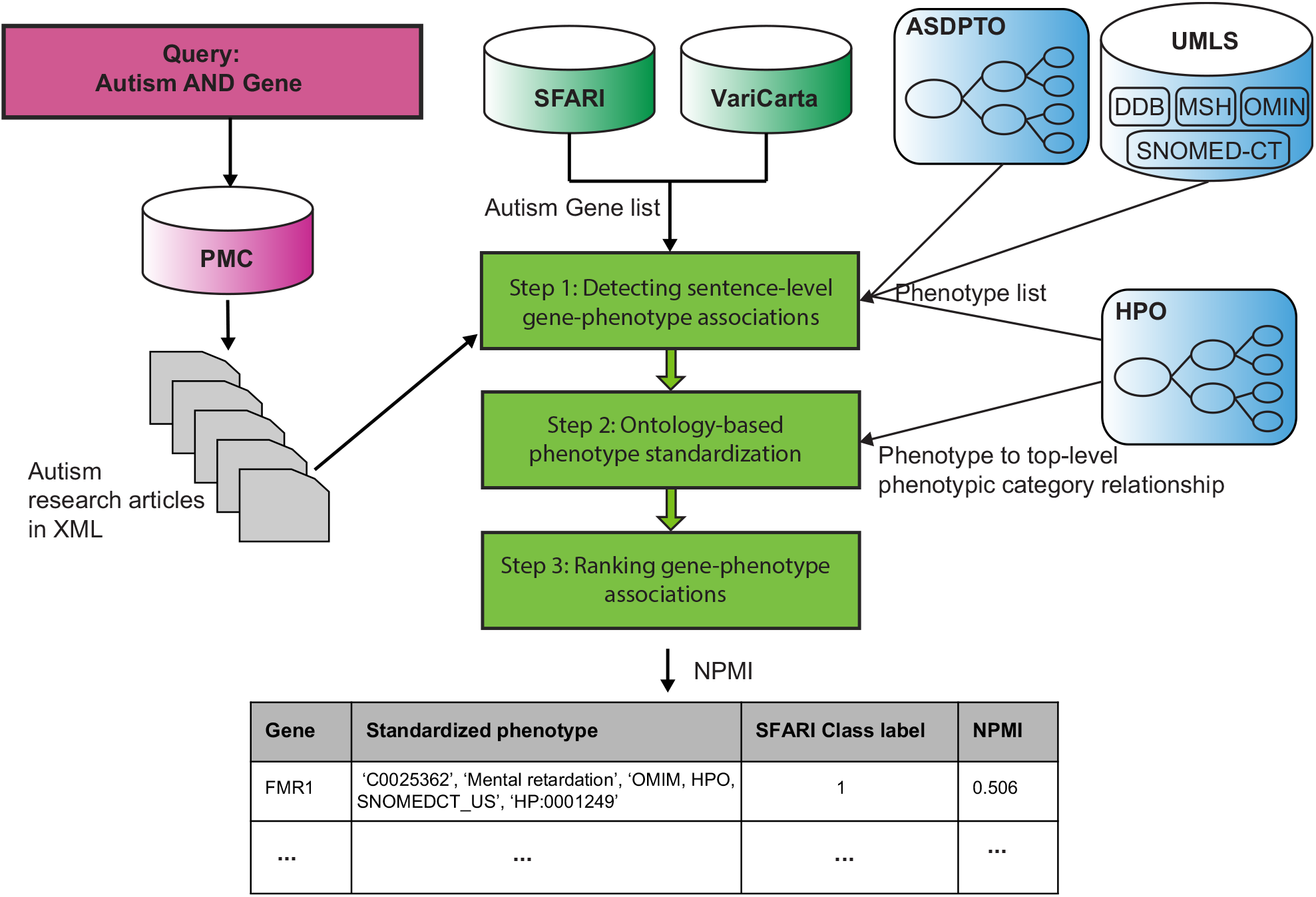
The overview pipeline of Autism_genepheno. For the standardized phenotype, it includes the unique UMLS concept ID, its preferred name in UMLS, its vocabulary source, and its corresponding HPO ID if it exists.

### Data Resources

We searched 15,070 autism research articles in PMC as our corpus. The search criteria were defined as including the keywords ‘Autism’ AND ‘Gene’ over the last 5 years, from 2015 to 2020.

We used 19,979 genes from VariCarta^20^, a comprehensive database of harmonized genomic variants found in autism spectrum disorder sequencing studies. Among 19,979 genes in VariCarta, we further identified 992 autism-associated genes from the SFARI database, and they were categorized into four classes (class 1: High Confidence, N= 194; class 2: Strong Candidate, N= 207; class 3: Suggestive Evidence, N= 507; class S: Syndromic, N= 84) based on their gene scores.

We constructed a phenotype list from two sources: Autism Spectrum Disorder Phenotype Ontology (ASDPTO) and Unified Medical Language System (UMLS)^21^. The phenotype list covered 284 concepts from ASDPTO; 33,384 concepts in Human Phenotype Ontology (HPO), 31,822 concepts in Online Mendelian Inheritance in Man (OMIM), 102 concepts in Diseases Database (DDB), 208,086 concepts in the Systematized Nomenclature of Medicine-Clinical Terms (SNOMED-CT, US edition) and 56,825 concepts in the Medical Subject Headings (MSH), respectively from UMLS. We extracted the above phenotype concepts from UMLS by filtering two data files MRCONSO.RFF and MRSTY.RFF from the UMLS Metathesaurus 2020AA. The file MRCONSO.RFF includes all concepts with their concept identifier, concept names, connected languages and source vocabularies. The file MRSTY.RFF provides a semantic type for each concept. The two layer filtering includes: 1) Filtering by source vocabularies, we added the concepts from MRCONSO.RFF into the phenotype list by selecting HPO, OMIM, DDB, SNOMED-CT and MSH as the source vocabularies. 2) Filtering by semantic types, we selected the concepts from the above five source vocabularies with their corresponding semantic types. We included concepts with semantic types of T047 (Disease or Syndrome), T048 (Mental or Behavioral Dysfunction) and T184 (Sign or Symptom) for the source vocabularies OMIM, DDB, SNOMED-CT and MSH. For HPO, we included all concepts to the phenotype list excluding concepts with semantic types of T045, T077, T079, T080, T082 or T169, etc.

### Detecting sentence-level gene-phenotype associations

We detected phenotype and gene information at the sentence-level from autism research articles in XML format with NLP methods. The step of detecting the sentence-level gene-phenotype associations in each article included four sub-steps: 1) process each article into sentence level with tokenizer, stop-words remover and lemmatizer; 2) tokenize and lemmatize all terms in the phenotype list that is constructed with ASDPTO and UMLS; 3) identify target sentence if at lease one tokenized phenotype term from the phenotype list and one gene symbol from the gene list are mentioned in the processed sentence with sub-string match; 4) extract all mentions of phenotype and gene in target sentence. Finally we detected the sentence-level gene-phenotype ASD association in the corpus.

### Ontology Based Phenotype Standardization

The mentions of phenotype identified in the corpus have multiple verbal expression, synonyms and lexical variants due to human language feature. We developed a source based method to standardize the mentions of phenotype in a consistent and systematic way. The source based method used the sources of the constructing phenotype list to standardize the mentions of phenotype with the most commonly preferred name in standard biomedical vocabularies. The Human Phenotype Ontology (HPO) is the highest priority source for a standard vocabulary of human phenotype. Other standard vocabularies were utilized as a complement to the HPO with the following descending priority order: ASDPTO, OMIM, DDB, SNOMED-CT and MSH. The standardization of phenotype with HPO further enabled us to group these identified phenotypes into 23 top-level phenotypic categories of phenotypic abnormality: Abnormal cellular phenotype, Abnormality of blood and blood-forming tissues, Abnormality of head or neck, Abnormality of limbs, Abnormality of metabolism/homeostasis, Abnormality of prenatal development or birth, Abnormality of the breast, Abnormality of the cardiovascular system, Abnormality of the digestive system, Abnormality of the ear, Abnormality of the endocrine system, Abnormality of the eye, Abnormality of the genitourinary system, Abnormality of the immune system, Abnormality of the integument, Abnormality of the musculoskeletal system, Abnormality of the nervous system, Abnormality of the respiratory system, Abnormality of the thoracic cavity, Abnormality of the voice, Constitutional symptom, Growth abnormality and Neoplasm. We used these top-level phenotypic categories to label autism-associated genes in the analysis that followed. If the standardized phenotype was not included in HPO, we defined its top-level phenotypic category as “NA”.

### Ranking gene-phenotype associations

To select high-confidence gene-phenotype associations, we ranked each association between a gene and phenotype using Normalized Pointwise Mutual Information (NPMI). NPMI is a measure of association between two terms^22, 23^. NPMI ranges from −1 to 1, with −1 indicating that the two terms never occurred together, 0 indicating that the terms occurrence independently, and 1 indicating that the terms always co-occurred together. In our study, the two terms are gene and phenotype. We calculate the NPMI between a gene G and a phenotype P as

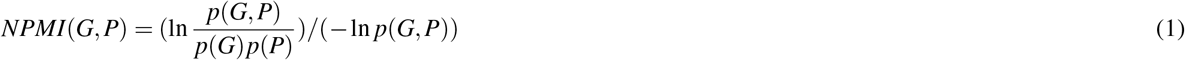

where *p*(*G*) and *p*(*P*) are the probability of gene and phenotype occurring in the corpus separately, and *p*(*G*, *P*) is the observed probability of gene and phenotype occurring in the same sentence. In our study, *p*(*G*) = *n*_*G*_/*n*_*tot*_, where *n*_*G*_ is the number of sentences containing the gene, and *n*_*tot*_ is the total number of sentences in our corpus; *p*(*P*) = *n*_*P*_/*n*_*tot*_, where *n*_*P*_ is the number of sentences mentioning the phenotype; *p*(*G*, *P*) = *n*_*G,P*_/*n*_*tot*_, where *n*_*G,P*_ is the number of sentences where the gene and phenotype co-occurs. Thus, the NPMI of a gene and a phenotype is calculated by the following formula:

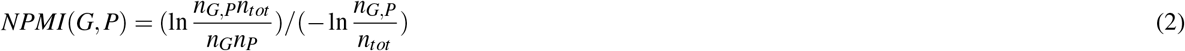

### Autism-associated gene classification by gene-phenotype associations

We constructed a gene-phenotype matrix in which each row represented a gene and each column represented a phenotype. Each entry in the matrix was set to “1” if the NPMI of the gene-phenotype association was larger than zero, and “0” if the NPMI of the gene-phenotype association was less than or equal to zero. We then performed dimensionality reduction through t-distributed stochastic neighbor embedding (t-SNE) and Kmeans clustering for the gene-phenotype matrix, to identify different clusters of autism-associated genes.

### Evaluation metrics of clustering results

We used four evaluation methods for clustering results: adjusted Rand index, Jaccard index, normalized mutual information, and purity score, to analyse the concordance between the phenotype’s top-level phenotypic category labels and the Kmeans clustering results on the two dimensions of the t-SNE results. Most of these four measurements vary from 0 to 1, with 1 indicating perfect match between them, except the adjusted Rand index which could yield negative values when concordance is less than expected by chance. The adjusted Rand index is an adjusted version of Rand’s statistic which is the probability that a randomly selected pair is classified in agreement. The Jaccard index is similar to Rand Index, but disregards the pairs of elements that are in different clusters for both clusterings. The normalized mutual information combines multiple clusterings into a single one without accessing the original features or algorithms that determine these clusterings. The purity score shows the rate of the total number of cells that are classified correctly.

### GO analysis

GO and Kyoto Encyclopedia of Genes and Genomes (KEGG) pathway enrichment analyses were conducted for each cluster of autism-associated genes from Kmeans clustering results through the Database for Annotation, Visualization and Integrated Discovery (DAVID) v6.8 (https://david.ncifcrf.gov/).

### Genetic interaction network graph

We constructed a genetic interaction network graph for SFARI genes in our database for the later analysis. The nodes/vertices in this network graph were SFARI genes and an edge was added to the graph if a certain pair of nodes shared a standardized phenotype (we used the global average NPMI value as the threshold for both genes sharing the same phenotype). If multiple edges existed between two nodes, these edges were compressed (combined) and the total number of edges were treated as the weight of the compressed edge.

We used betweenness centrality to rank SFARI genes. The betweenness centrality is widely used in graph analysis as a measure of centrality based on shortest paths^24, 25^. It is used to quantify the amount of influence a node (SFARI gene in our genetic interaction graph) has over the flow of information in a graph. The betweenness centrality calculates the shortest (weighted) path between every pair of nodes in a connected graph. Nodes (SFARI genes) will receive a higher betweenness centrality score if it most frequently lie on these shortest paths and serve as a bridge among all nodes in the graph. The betweenness centrality *C*_*B*_(*v*) for a node *v* is defined as below:

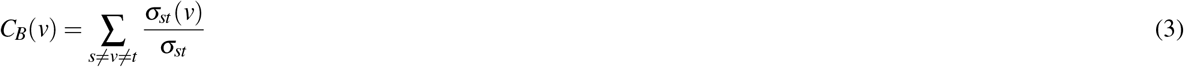

where *σ*_*st*_ is the total number of shortest paths from node *s* to node *t* and *σ*_*st*_ (*v*) is the number of those paths that pass through *v*.

## Results

We applied Autism_genepheno to five years of literature that included the keywords autism and gene (N = 15,070), and extracted a total of 71,558 gene–phenotype associations from 6,892 autism-associated genes and 5,493 standardized phenotypes (2742 from HPO, 46 from ASDPTO, 119 from OMIM,2 from DDB, 1418 from SNOMED-CT, US edition and 1133 from MSH) (Methods section). Before analyzing phenotypic associations, we tested how well genes identified with this process corresponded with those identified in a well-known database of autism genomic data, SFARI Gene 3.0, which includes approximately 1,000 genes associated with autism risk. SFARI genes are categorized into four classes, associated with ASD risk at different confidence levels (Methods section). Our results identified gene-phenotype associations for 751 SFARI genes, of which 172 were SFARI genes labeled as class 1 (conferring highest autism risk), 154 were SFARI genes labeled as class 2, 349 were SFARI genes labeled as class 3, and 76 were SFARI genes labeled as class S. The rest 6,139 genes were not included in the SFARI autism gene database, but were included in the VariCarta autism gene database. In this paper, we labeled the rest of the genes as “NA”.

### Top mentioned genes and phenotypes in ASD

For a general description of all genes and associations, we simply applied *NPMI* > 0 and *n*_*G,P*_ > 5 as thresholds to filter out low-confidence gene-phenotype associations. The result showed that SFARI genes accounted for 29.8% of the total percentages of genes extracted by Austim_genepheno and SFARI gene-phenotype accounted for 44.9% of the total gene-phenotype associations. Class 1 SFARI genes in particular (red bars in Fig 1A) made up 10.6% of the genes but accounted for more than twice as many associations (24.5%). These statistics indicated that the highest-risk SFARI genes were most frequently mentioned in the context of ASD phenotype associations.

To investigate whether the most frequently mentioned genes in our data sources matched the confidence pattern observed in SFARI Gene, we plotted the top 30 genes in terms of number of papers referencing each gene. The plot in Figure 2B shows that more than half (18) of the genes, including the top 9 genes with the highest frequencies were SFARI genes labeled with class 1 (red bars). SFARI genes labeled with class 2 (blue bars) made up the second largest group, followed by genes labeled with class 3 (green bars) and gene labeled with class S (yellow bars). This result confirmed that frequency of gene reference in the literature alone is well predictive of risk in ASD. Indeed, the SFARI Gene system integrates gene frequencies from literature mining into their scoring model.

**Figure 2.**
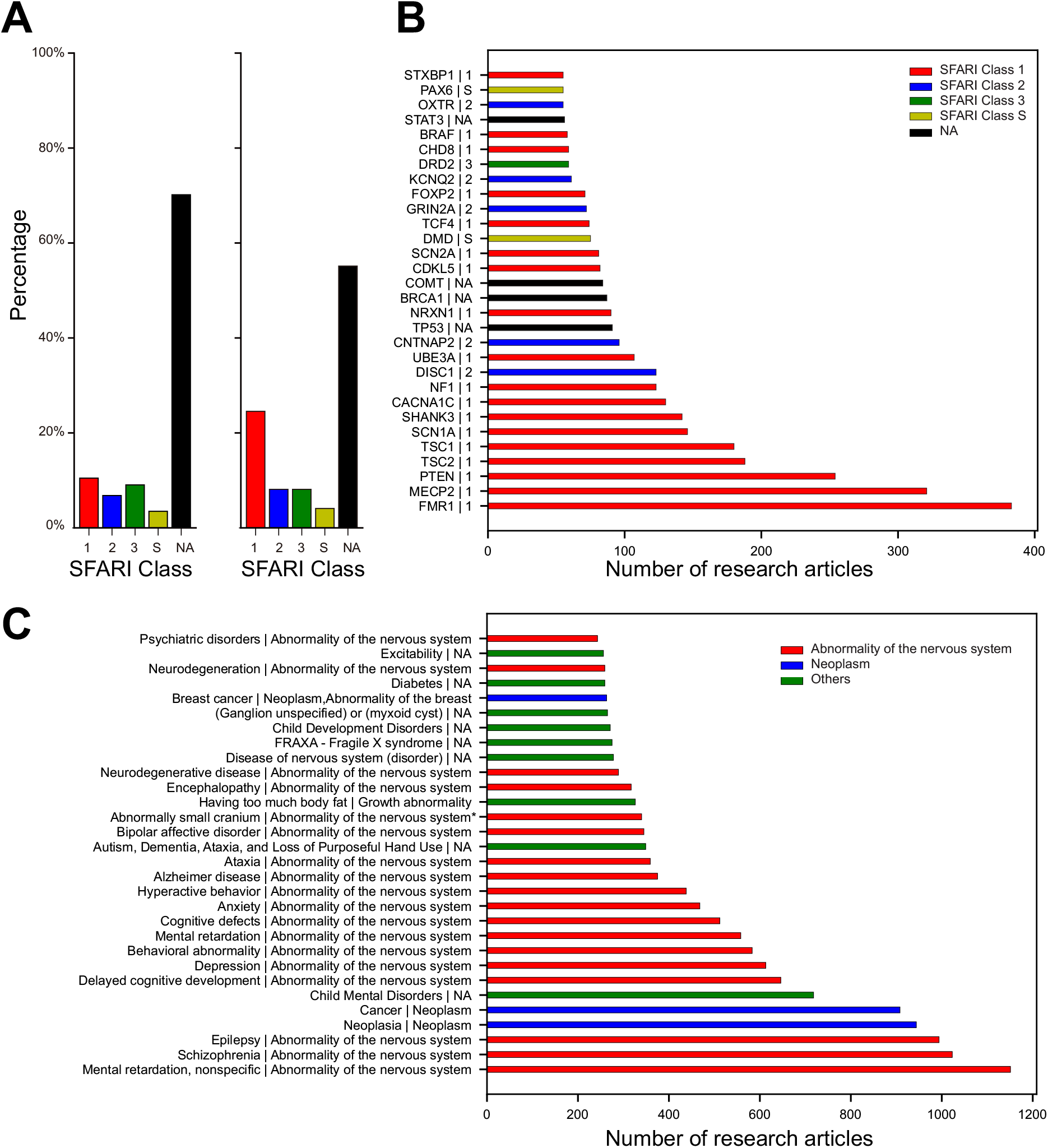
Top mentioned autism-associated genes correspond to the classes of SFARI genes. A. The percentage distribution of different classes of SFARI genes and “NA” genes in our data resource (left panel). The percentage distribution of different classes of SFARI genes associated standardized phenotypes and “NA” genes associated standardized phenotypes in our data resource (right panel). B. Top 30 mentioned autism-associated genes. C. Top 30 mentioned autism-associated standardized phenotypes. The standardized phenotype and top-level phenotypic category are separated by “|” here for each term. Top-level phenotypic category “NA” means the standardized phenotype is not included in HPO.

This analysis also identified four genes (TP53, BRCA1, COMT and STAT3) not included in the most recent SFARI gene list, and thus labeled as “NA” in our plot (black bars). We were interested to investigate whether this occurance was spurious, or reflected a hihterto unnoticed autism association. In table 1, we randomly selected eight SFARI genes (two genes for each class label, and three high-confidence standardized phenotypes with highest NPMI values where *n*_*G,P*_ > 5 for each gene, respectively), and four additional “NA” genes. We evaluated all these gene-phenotype associations through the OMIM database. The TP53 gene encodes a protein called tumor protein p53 (or p53) that acts as a tumor suppressor to regulate cell division by keeping cells from growing and dividing in an uncontrolled way. The top three phenotypes extracted by Austim_genepheno for TP53 were Li-Fraumeni Syndrome, neoplasia, and cancer (Table 1). The top three phenotypes for the BRCA1 gene were ovarian cancer, breast cancer and cancer, and the top three phenotypes for the STAT3 were prostate cancer, retinoblastoma, and Buckley syndrome. The phenotypes from the three genes suggested that biological processes related to cancer could play an important role in ASD that has been missed by the SFARI database. Recent studies have also shown that autistic brains share gene expression and biological pathway abnormalities with cancer^26–28^. The top three phenotypes for COMT gene were 22q11 microdeletion with velocardiofacial syndrome phenotype, schizophrenia, and aggression. Emerging studies also suggest that there are both clinical and biological links between autism and schizophrenia^29–31^.

**Table 1.** Gene-phenotype association for eight SFARI genes and four “NA” genes. Each gene is associated with three standardized phenotypes with the highest NPMI values.

| Gene | Standardized Phenotype | SFARI Class Label | NPMI |
| --- | --- | --- | --- |
| FMR1 | FXTAS Fragile X Tremor Ataxia Syndrome | 1 | 0.514 |
| FMR1 | Mental retardation | 1 | 0.506 |
| FMR1 | FRAXA - Fragile X syndrome | 1 | 0.497 |
| MECP2 | Lubs X-linked intellectual disability syndrome | 1 | 0.681 |
| MECP2 | Autism, Dementia, Ataxia, and Loss of Purposeful Hand Use | 1 | 0.572 |
| MECP2 | Neonatal encephalopathy | 1 | 0.469 |
| CNTNAP2 | CDFE (cortical dysplasia focal epilepsy) syndrome | 2 | 0.620 |
| CNTNAP2 | Language impairment | 2 | 0.399 |
| CNTNAP2 | Specific Language Disorder | 2 | 0.381 |
| GRIN2A | Acquired Aphasia with Convulsive Disorder | 2 | 0.596 |
| GRIN2A | Epilepsy, Rolandic | 2 | 0.560 |
| GRIN2A | Epilepsies, Partial | 2 | 0.445 |
| DRD2 | Alcoholism | 3 | 0.364 |
| DRD2 | Addictive behavior | 3 | 0.276 |
| DRD2 | Increased body weight | 3 | 0.233 |
| BRCA2 | Ovarian cancer | 3 | 0.501 |
| BRCA2 | Fanconi Anemia | 3 | 0.439 |
| BRCA2 | Breast cancer | 3 | 0.436 |
| CACNA1A | Familial Hemiplegic Migraine | S | 0.661 |
| CACNA1A | Ataxia 6s, Spinocerebellar | S | 0.657 |
| CACNA1A | Hemiplegic migraine | S | 0.654 |
| DMD | Benign Duchenne muscular dystrophy | S | 0.673 |
| DMD | Muscular dystrophy | S | 0.607 |
| DMD | BMD - Becker muscular dystrophy | S | 0.592 |
| COMT | 22q11 microdeletion with velocardiofacial syndrome phenotype | NA | 0.274 |
| COMT | Schizophrenia | NA | 0.184 |
| COMT | Aggression | NA | 0.176 |
| TP53 | Li-Fraumeni Syndrome | NA | 0.579 |
| TP53 | Neoplasia | NA | 0.317 |
| TP53 | Cancer | NA | 0.293 |
| BRCA1 | Ovarian cancer | NA | 0.550 |
| BRCA1 | Breast cancer | NA | 0.465 |
| BRCA1 | Cancer | NA | 0.339 |
| STAT3 | Prostate cancer | NA | 0.279 |
| STAT3 | Retinoblastoma | NA | 0.410 |
| STAT3 | Buckley Syndrome | NA | 0.520 |

We also wished to test whether the most frequently mentioned standardized phenotypes in our data sources corresponded well to autism pathologies. We therefore plotted the top 30 mentioned standardized phenotypes and their respective top-level phenotypic categories (Figure 2C). The two most common mentioned top-level phenotypic categories were abnormality of the nervous system and neoplasm (Figure 2C, red and blue bars). Among other top-level phenotypic categories (Figure 2C, green bars), the most common standardized phenotypes ordered by frequency were: 1) Child mental disorders, 2) Autism, dementia, ataxia, and loss of purposeful hand use, 3) Having too much body fat, 4) Disease of nervous system (disorder), 5) FRAXA - Fragile X syndrome, 6) Child development disorders, 7) Ganglion unspecified or myxoid cyst, 8) Diabetes, and 9) Excitability. We conclude that our pipeline identified both genes and phenotypes highly relevant for autism.

### Evaluation of gene-phenotype associations from Autism_genepheno

To first evaluate all gene-phenotype associations extracted by the pipeline, we manually curated 50 papers to serve as a gold standard and built a metric to evaluate our results (see details in Supplementary file). We used precision and recall metrics at the sentence, gene and phenotype level to evaluate the pipeline’s performance. The results are shown in table 2. We extracted a total of 71,558 gene–phenotype associations. At the gene level, the recall and precision were approximately 91.4% and 74.4%, respectively. At the phenotype level, the recall and precision were approximately 77.1% and 98.7%, respectively. As we expected, these evaluation results suggested that genes were straightforward to detect but with a higher false positive rate, and phenotypes were intricate to detect but with a lower false positive rate. At the sentence level, recall and precision were approximately 80.3% and 71.9%, respectively.

**Table 2.** Gene-phenotype associations evaluation from gene, phenotype and sentence levels by manual annotations from 50 autism research articles.

|  | Gene level | Phenotype level | Sentence level |
| --- | --- | --- | --- |
| Benchmark | 175 | 193 | 137 |
| True Positive | 160 | 148 | 110 |
| False Positive | 55 | 2 | 43 |
| False Negative | 15 | 44 | 27 |
| Precision | 74.4% | 98.7% | 71.9% |
| Recall | 91.4% | 77.1% | 80.3% |

To further evaluate the gene–phenotype associations with positive NPMI values, we used a reference annotation database from the HPO (https://hpo.jax.org/app/download/annotation). A total of 2,913 genes identified by Autism_genepheno were present in HPO. If there was at least one gene-phenotype association found both in the reference database and our autism gene-phenotype association database for each gene, we defined the autism-associated gene as a true positive (TP). A total of 94.4% (136/144) SFARI genes labeled as class 1 were evaluated as TPs, followed by 82.9% (68/82) for SFARI genes labeled as class 2, 80.1% (129/161) SFARI genes labeled as class 3, 83.3% (60/72) SFARI genes labeled as class S, and 62.4% (1531/2454) “NA” genes, not included in SFARI (Figure 3). This evaluation result showed the SFARI class 1 genes conferring the highest autism risk had the highest true positive rate, followed by SFARI class 2 and 3 genes. The result indicates that Autism_genepheno successfully extracted gene-phenotype associations, particularly for genes with high autism risk.

**Figure 3.**
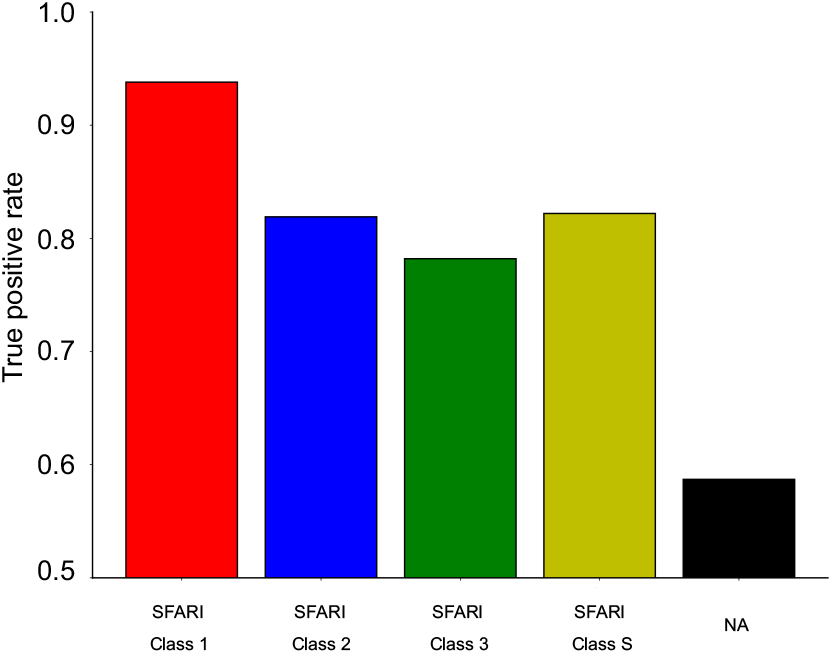
Gene-phenotype evaluation rate through HPO for all classes of SFARI genes and “NA” genes which are not included in the SFARI database but in the VariCarta database.

### Autism-associated gene classification by top-level phenotypic categories of standardized phenotypes

To investigate whether we could classify autism-associated genes extracted from Autism_genepheno based on gene-phenotype associations, we first generated a matrix from all the gene-phenotype associations, where each row represented a gene and each column represented a phenotype. Each entry in the matrix was set to “1” if the NPMI of the gene-phenotype association was larger than zero, and “0” if the NPMI of the gene-phenotype association was less than or equal to zero. We then applied t-SNE to this gene-phenotype matrix (see Methods Section) and labeled genes based on their corresponding top-level phenotypic category (Fig. 3A). Since gene-to-phenotype mapping was not one-to-one but one-to-many, we selected the standardized phenotype with the highest NPMI for each gene and then obtained its top-level phenotypic category of this phenotype. Thus we only kept 3,683 autism-associated genes whose phenotypes with the highest NPMI were present in HPO since they could then be traced back to their corresponding top-level phenotypic categories. We then performed Kmeans clustering and produced seven gene clusters (Figure 4B). The top and bottom gene clusters matched in the two scatter plots (4A vs. 4B), which means that the strongest phenotypes associated with each gene (standardized phenotypes with the highest NPMI) was one of the main factors distinguishing different gene clusters. To quantify the concordance result of these two clusters from two different clustering and labeling systems, we evaluated it by four different metrics: adjusted Rand index, Jaccard index, normalized mutual information, and purity (Methods section). The adjusted Rand index, Jaccard index, normalized mutual information, and purity score achieved 0.45, 0.53, 0.53 and 0.67, respectively.

**Figure 4.**
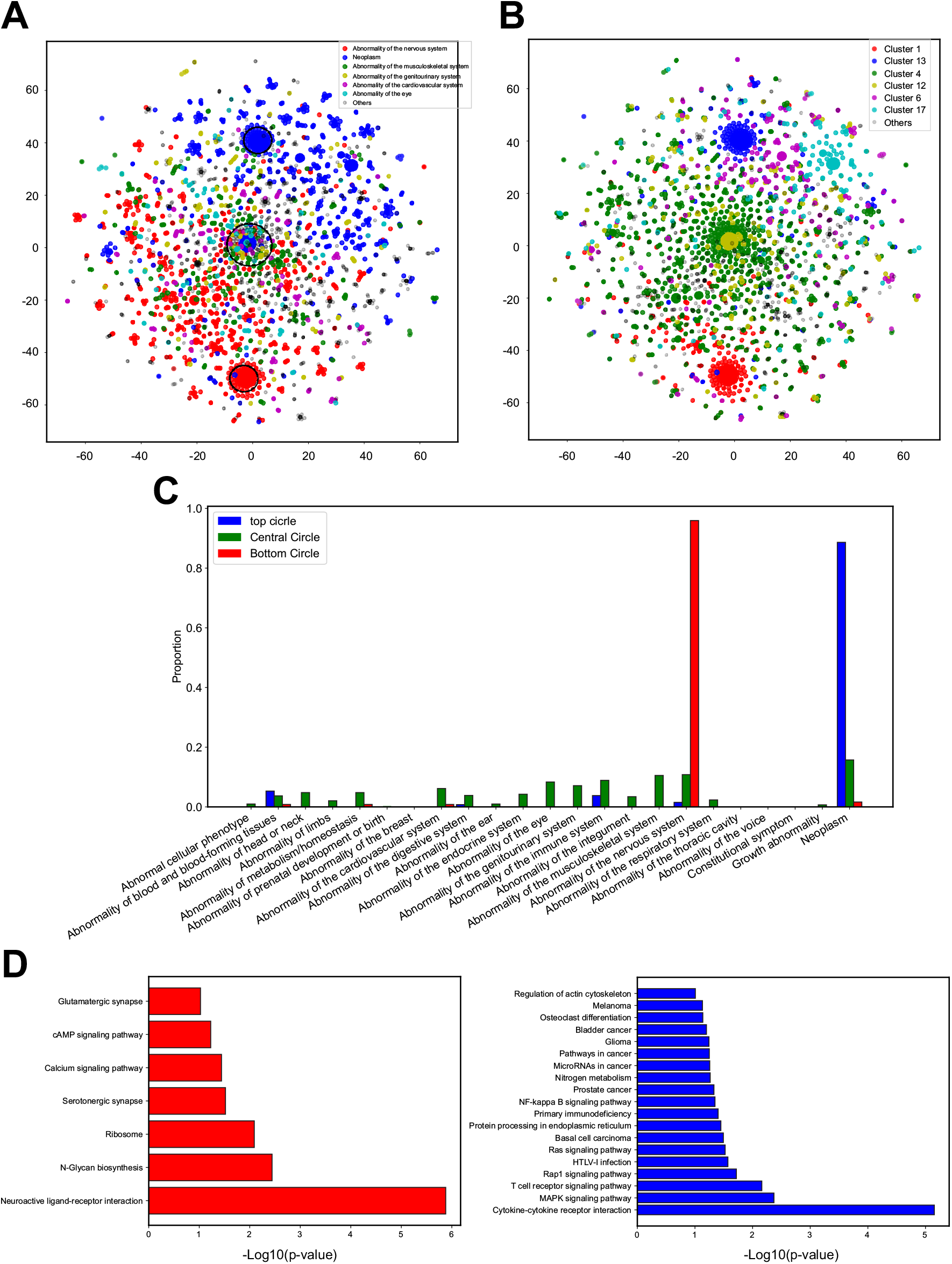
Autism-associated gene classification through top-level phenotypic categories and Kmeans clustering. A. The t-SNE plot of all autism-associated genes labeled by top-level phenotypic categories. B. The t-SNE plot of all autism-associated genes labeled by Kmeans clustering results. C. Distribution of top-level phenotypic categories for top, central and bottom gene clusters marked in black circles from A. D. The GO analysis for cluster 1 (red cluster) and 13 (blue cluster) from Kmeans clustering.

As mentioned above, the top gene cluster (genes within the black circle of the top gene group) corresponded to the neoplasm top-level phenotypic category, and the bottom gene cluster (genes within the black circle of the bottom gene group) corresponded to the abnormality of the nervous system top-level phenotypic category. We then selected genes (dots of seven different colors) within the central circle as the third gene group (Figure 4B). These genes were classified into many different top-level phenotypic categories: abnormality of the nervous system, neoplasm, abnormality of the musculoskeletal system, abnormality of the genitourinary system, abnormality of the cardiovascular system, abnormality of the eye, and others (Figure 4A). Even though the gene-to-phenotype mapping was one-to-many and we only chose standardized phenotypes with the highest NPMI to annotate each gene, each gene of the top and bottom groups was more likely to be associated with only one standardized phenotype. On the other hand, genes of the central group were more likely to be associated with many phenotypes. To demonstrate this, we plotted a percentage distribution of all phenotypes of genes for all three gene groups we marked in the circles. As expected, the phenotypes for the top group were dominated by the top-level phenotypic category of neoplasm (blue bars), and the phenotypes for the bottom group were dominated by the top-level phenotypic category of abnormality of the nervous system (red bars). However, the phenotypes of the central group were evenly distributed among several different upper classes (green bars). These results revealed the phenotypic pattern of autism-associated genes.

Although our analysis produced well defined clusters of genes based on associations retrieved from processing the research literature with Autism_genepheno, we wanted to ensure that these corresponded with known gene functions. We thus, performed GO analysis for genes within two clusters (Cluster 1 and 13) obtained via the Kmeans clustering results (Figure 4B). The GO results indicated that cluster 1 genes were most involved in neuroactive ligand-receptor interaction and several other components and pathways (N-Glycan biosynthesis, ribosome, serotonergic synapse, calcium signaling pathway, cAMP signaling pathway, and glutamatergic synapse). In every instance, the gene ontology revealed functions or pathways present in neurons, whose malfunctions were associated with the top-level phenotypic category of abnormality of the nervous system (Figure 4D, left panel). For cluster 13, all the GO terms (such as cytokine-cytokine receptor interaction, MAPK signaling pathway, T cell receptor signaling pathway, Ras signaling pathway, prostate cancer, glioma, and melanoma) were relevant to the top-level phenotypic category of neoplasm (Figure 4D, right panel). These results independently confirmed that Autism_genepheno recovered phenotypes correctly.

### The unique top-level phenotypic category profiles of SFARI genes and the genetic interaction network

We then wished to test if autism risk, as indicated based on SFARI gene class, correlated strongly with specific phenotypes, or top-level phenotypic categories. We thus plotted the spatial distribution of SFARI genes of different classes within the pattern of autism-associated genes identified based on phenotype. We labeled each gene from the SFARI Gene database based on its risk score, and labeled the gene as “NA” if it did not belong to the SFARI Gene database. Figure5 shows that the 283 SFARI genes out of total 3,683 genes were spread widely in the t-SNE plot, but did not lie in the top, bottom and central clusters which were mainly dominated by the other genes labeled as “NA” (Figure 5A). This suggested that the phenotypic pattern of SFARI genes was highly variable and autism risk did not correlate strongly with specific phenotype clusters. To investigate the difference of phenotypic pattern for different classes of SFARI genes (class 1: 45, 2:77, 3:133 and S:28), we calculated the proportion of each class of SFARI genes belonging to each top-level phenotypic category (Figure 5B). To calculate this, for each gene, we selected the standardized phenotype with the highest NPMI and obtained its top-level phenotypic category. This result indicated that each class of SFARI genes had some unique phenotypic pattern. First, the SFARI genes associated with high confidence level autism risk were more involved in 3 top-level phenotypic categories: abnormality of blood and blood-forming tissues, abnormality of the integument, and abnormality of head or neck. Secondly, SFARI genes labeled as class 2 in terms of autism risk were more associated with 7 top-level phenotypic categories: abnormality of limbs, abnormality of metabolism/homeostasis, abnormality of the cardiovascular system, abnormality of the digestive system, abnormality of the musculoskeletal system, abnormality of the ear, and abnormality of head or neck. Thirdly, SFARI genes labeled as class 3 in terms of autism risk were more involved in 14 top-level phenotypic categories: abnormal cellular phenotype, abnormality of blood and blood-forming tissues, abnormality of limbs, abnormality of the digestive system, abnormality of the ear, abnormality of the endocrine system, abnormality of the genitourinary system, abnormality of the immune system, abnormality of the musculoskeletal system, abnormality of the nervous system, abnormality of the respiratory system, constitutional symptom, growth abnormality and neoplasm. Lastly, the SFARI genes labeled as syndromic only contributed to a small proportion of top-level phenotypic categories, such as abnormality of metabolism/homeostasis, abnormality of the endocrine system, and abnormality of the integument.

**Figure 5.**
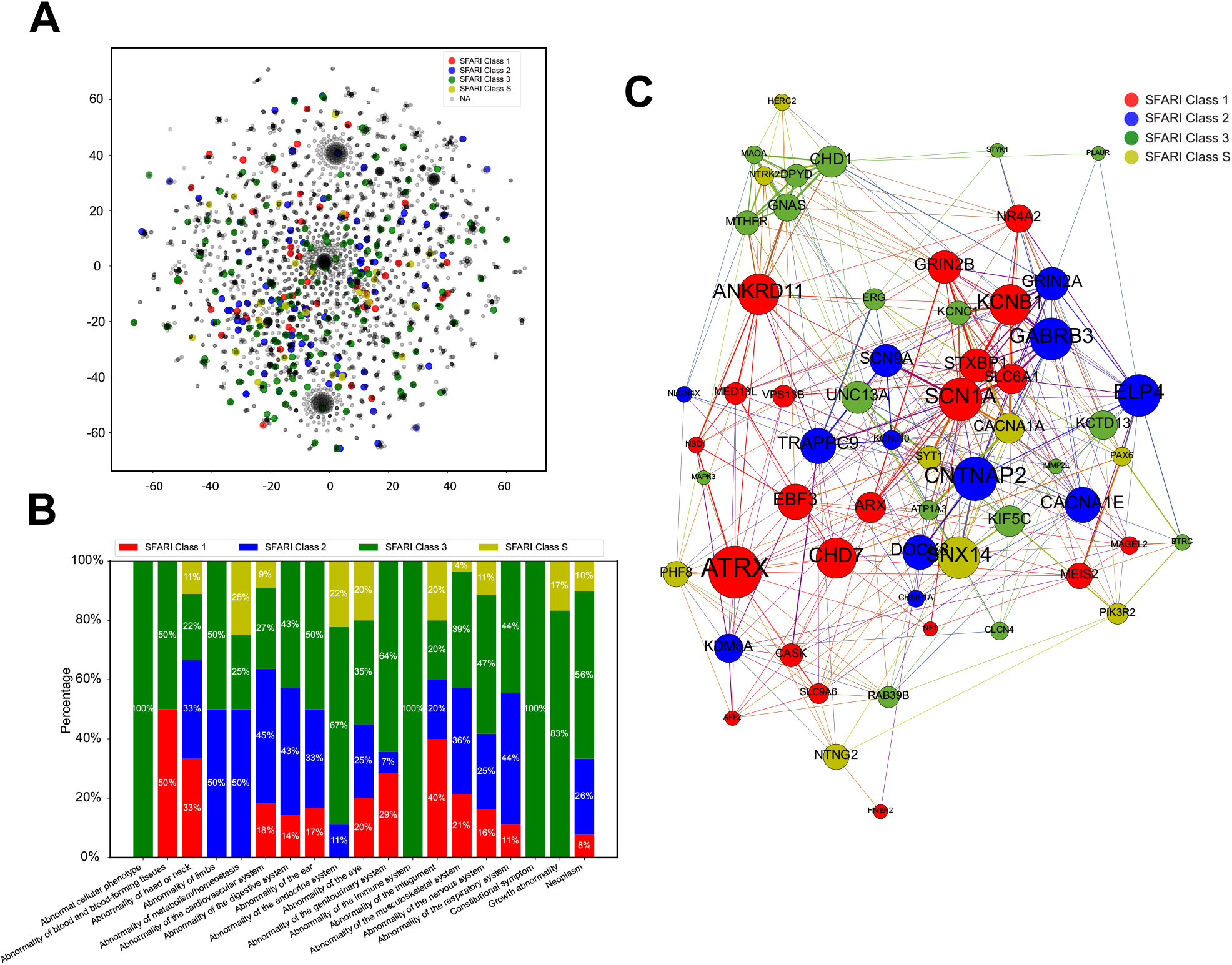
The unique phenotypic profiles of four classes of SFARI genes. A. The spatial distribution of SFARI genes on the t-SNE plot. B. The proportion of each class of SFARI genes belongs to each top-level phenotypic category. Red: SFARI class 1, Blue: SFARI class 2, Green: SFARI Class 3, and Yellow: SFARI Class S. C. The genetic interaction network graph of those top 10% SFARI genes with highest betweenness centrality scores.

As we saw above, SFARI genes were widely spread in the t-SNE plot and demonstrated a unique and wide spectrum of phenotypes for different classes. To further rank SFARI genes and detect the group of autism-associated genes playing important roles in connecting different genes together based on their phenotypic profiles, we constructed a genetic interaction network graph (see Methods section). We calculated the betweenness centrality scores for all SFARI genes and selected the top 10% of SFARI genes (N = 60) with the highest betweenness centrality scores (Supplementary table). The genetic interaction network graph for these 60 genes is shown in Figure 5C, with the size of each node corresponding to its betweenness centrality score. The top 10% of nodes with the highest betweenness centrality scores included 15.9% (21/132) SFARI class 1, 8.1% (12/148) class 2, 6.3% (18/287) class 3, and 13.8% (9/65) class S genes.

## Discussion and Conclusion

We introduce here an automatic text mining pipeline to extract gene-phenotype associations from the autism research literature. At present, gold-standard databases require substantial manual curation, however as the publication rate accelerates this may not be feasible in the future. Our approach integrates a comprehensive phenotype ontology list from the Autism Spectrum Disorder Phenotype Ontology and Unified Medical Language System, and a comprehensive autism-associated gene list from the VariCarta and the SFARI databases. We generated 71,558 gene-phenotype associations for a total of 6,892 autism-associated genes. All our data resources are freely available and will be updated frequently as more autism related articles are published. Our pipeline thus provides a resource that researchers can use at any time, to obtain the most up-to-date understanding of phenotypes associated with genes implicated in autism.

To evaluate the performance of our pipeline, we manually curated 50 articles and calculated the recall and precision rates for sentence-level extraction of gene-phenotype associations (80.3% for recall and 71.9% for precision). We then used a reference database to further evaluate the performance. SFARI class 1, 2, 3, S genes and NA genes achieved true positive rates of 94.4%, 82.9%, 80.1%, 83.3%, and 62.4%, respectively. This evaluation result corresponded to the SFARI gene class/scoring system in which SFARI class 1 genes conferred the highest autism risk, followed by SFARI class 2, 3 genes. In GO analysis for specific clusters of autism-associated genes, we further confirmed that phenotypes extracted by the pipeline were consistent with biological functions and pathways.

Since the same gene could lead to different disease symptoms, Autism_genepheno generated a specialized database of autism-associated gene-phenotype associations, which were specifically mentioned in autism research, rather than a generalized database of gene-phenotype associations for all genetic studies. To investigate the phenotypic pattern of each autism-associated gene, our pipeline provides a more autism-oriented database. We were able to demonstrate the unique and wide spectrum of phenotypes for SFARI genes from the top-level phenotyic categories, and ranked the 10% SFARI genes with the most influential roles in the genetic interaction network. These results could provide a unique resource for the diagnosis and treatment for ASD.

Our pipeline has some limitations. Our evaluation showed the gene-phenotype associations results contained both false positives and false negatives. False positives are caused by the following two reasons. First, due to our pipeline mistakenly identifying abbreviations as gene symbols. For instance, the pipeline identified ‘CARS’, the abbreviation of Childhood Autism Rating Scale, as a gene which encodes cysteinyl-tRNA synthetase (also an alias of gene CARS1). Secondly, the pipeline detected phenotype terms constructing non-phenotype noun phrases in the sentence. For example ‘cancer’ in the noun phrase ‘National Cancer Center’ was detected as the synonym of phenotypic abnormality ‘Neoplasm’ in HPO. Most false negatives were due to the inability to identify all possible phenotype expressions in the human language. Other false negatives come from ambiguity in processing gene symbols in non-normalized written forms. Our pipeline currently detects gene-phenotype associations in sentence-level by their co-occurrence and ranked their associations by NPMI only. The pipeline did not capture any linguistic relations between genes and phenotypes co-occurring in sentence-level, such as verbs indicating affirmation or negation. In future work, we suggest to rank gene-phenotype associations in sentence-level by capturing and quantifying their semantic relationships, such as causal relationships, declining relationships, and reinforcing relationships.

In the future, we also expect to adapt our pipeline to other neurological diseases, such as Alzheimer’s disease and Schizophrenia. Our approach can thus provide a comprehensive database of gene-phenotype associations for all neurological diseases to study the phenotypic pattern of each gene in different neurological diseases and their interactions.

## Supporting information

Autism_genepheno_SI

Autism_genepheno_SupplementaryTable

## Acknowledgments

The authors thank the anonymous reviewers for their valuable suggestions. This work is supported by Vanderbilt University Development Funds (Grant No. FF_300033).

## Author contributions

X.Z. and Y.Z. conceived and led this work. S.L, Z.G., Y.Z. and X.Z. designed the model and implemented the Autism_genepheno pipeline. S.L, Z.G., J.B.I., Y.H., Y.Z. and X.Z. led the data analysis. S.L, Z.G., Y.Z. and X.Z. wrote the paper with feedback from J.B.I. and Y.H.

