## Supplementary material for "Autism_genepheno: Text mining of gene-phenotype associations reveals new phenotypic profiles of autism-associated genes": Autism_genepheno_SI

### Rules of manual annotations

We manually curated XXX articles and recorded all sentences including genes and phenotypes which are marked in green. Examples of manual annotation are shown as below:

A. *"Among these genes, TBR1—a putative transcription factor (TF) —is highly expressed in glutamatergic early-born cortical neurons; it dictates the expression of other risk genes, controls cortical development, and is implicated in intellectual disability. "*
   Output: All found (TF is mistakenly found).
B. *"Following TBR1 dictationnd play, FOXP2 is also related to some severe speech-language disorders as a role in cortical neurogenesis."*
   Output: Sentence missed (No phenotype severe speech-language disorders in the list).
C. *"As evidence of its involvement in neurodevelopmental disorders, UBE3A is the causative gene of Angelman syndrome, a disorder resulting in the majority of cases from deletion of maternal 15q11.2-q13.1 and characterized by phenotypic overlap with Dup15q syndrome."*
   Output: Sentence found (Phenotype Dup15q syndrome missed)

**There were three types of annotation output:**

I. All found: genes and phenotypes were perfectly found by the pipeline.
II. Sentence missed: None of gene and phenotype in the sentence were detected by the pipeline.
III. Sentence found: The sentence including at least one gene and one phenotype was found by the pipeline, but not all of genes or phenotypes were detected.

For type II and III, the missed or detected gene or phenotype was provided in the parenthesis.

**Explanations of evaluation metrics:**

We used precision and recall metrics at sentence level, gene level and phenotype level to evaluate the pipeline's performance. The output of manual annotation served as the golden standard.

Precision = TP/ (TP + FP)

Recall = TP/(TP + FN)

(1) At sentence level:

True Positive (TP): The number of sentences that are both found in manual annotations and results of the pipeline.

False Positive (FP): The number of sentences that are found in results of the pipeline but not annotated.

False Negative (FN): The number of sentences that are manually annotated but not found by the pipeline.

(2) At genotype level:

True Positive (TP): The number of genotypes that are both found in manual annotations and results of the pipeline.

False Positive (FP): The number of genotypes that are extracted in results of the pipeline but not annotated.

False Negative (FN): The number of genotypes that are manually annotated but not found by the pipeline.

Each genotype in one sentence is only counted once no matter how many times it appears.

(3) At phenotype level:

True Positive (TP): The number of phenotypes that are both found in manual annotations and results of the pipeline.

False Positive (FP): The number of phenotypes that are extracted in results of the pipeline but not annotated.
False Negative (FN): The number of phenotypes that are manually annotated but not found by the pipeline.
Each phenotype in one sentence was only counted once no matter how many times it appeared.

To calculate FN at gene level and phenotype level, we only counted the missed gene or phenotype for the type II of annotation output-- 'Sentence missed'.
In the example B of manual annotation given above, only the phenotype 'severe speech-language disorders' was counted for FN at phenotype level while the gene 'TBR1' and 'FOXP2' were not counted for FN at gene level because we only missed the phenotype 'severe speech-language disorders'.

The most probable reason for most situations where the sentences were missed or where partial phenotypes are missed is that the missed phenotypes are not in the list generated from ASDPTO and UMLS.

**More example sentences:**

De novo CTNNB1 mutations have been reported in individuals with ASD, intellectual disability, microcephaly, motor delay, and speech impairment [36, 39, 61–63].
All found

These findings suggest that loss of Auts2 alters mouse vocal communication, which may underlie the pathology for communication disorders in patients with ASD with AUTS2 mutations.Figure 7Deficits in Vocal Communication in Adult Auts2del8 Mutant Mice(A) Representative spectrograms of USV during the courtship behaviors.
All found

Mutations in the NRXN1 gene have been implicated in a variety of conditions including autism, schizophrenia, and nicotine dependence (Ching et al., 2010).
All found

These variants prevent phosphorylation of DIXDC1 isoform 1, causing impairment to dendrite and spine growth [79].
Sentence missed (phenotype impairment to dendrite and spine growth missed).

More speculatively, SPRY3 deregulation could explain a reported, although currently unconfirmed, lung branching abnormality in autism (69).
Sentence missed (phenotype lung branching abnormality missed).

CTNNB1 haploinsufficiency has been found to cause neuronal loss, craniofacial anomalies, and hair follicle defects in both humans and mice [64].
Sentence found(Only phenotype neuronal loss found)

Previous studies have reported that mothers with NLRP5 mutations had offspring with characteristic clinical features and disorders, such as fertility impairment, infertility, idiopathic developmental delay and autism.
Sentence found (fertility impairment missed)

Conditional ablation of β-catenin in the dorsal neural folds of mouse embryos represses the expression of Pax3 and Cdx2 at the dorsal posterior neuropore and leads to a decreased expression of the WNT/β-catenin signaling target genes T, Tbx6, and Fgf8 at the tail bud, resulting in spina bifida aperta, caudal axis bending, and tail truncation [65].

Sentence found (Tbx6 and Fgf8 missed, only phenotype spina bifida found, caudal axis bending, tail truncation missed)

False positive sentences from the JSON outputs of Autism_genepheno:


"Sentence2": {
"Content": "APC The tumor suppressor APC is a key component of the β-catenin-destruction complex [56].",
"Gene": [
"APC"
],
"Original phenotype": [
"tumor"
],
"Standardized phenotype": [
"C0027651: Neoplasia(from HPO)"
]
},

"Sentence1": {
"Content": "GRK5, known as GPRK2 in flies, is a kinase that has been reported to be differentially methylated in schizophrenia studies [23, 32].",
"Gene": [
"GRK5"
],
"Original phenotype": [
"schizophrenia"
],
"Standardized phenotype": [
"C0036341: Schizophrenia(from MSH, OMIM, HPO, SNOMEDCT_US)"
]
}
}
